# Glycomic analysis reveals a conserved response to bacterial sepsis induced by different bacterial pathogens

**DOI:** 10.1101/2020.12.11.421610

**Authors:** Daniel W. Heindel, Peter V. Aziz, Shuhui Chen, Jamey D. Marth, Lara K. Mahal

## Abstract

Sepsis is an extreme inflammatory response to infection (bacterial, viral, fungal) that occurs in the bloodstream and causes damage throughout the body. Currently, there are few diagnostic biomarkers of sepsis and new effective treatments have not been developed. There is a clear need to study the molecular underpinnings of this disease. Glycosylation is known to play a role in immunity and inflammation, but the role of glycans in sepsis is not well defined. Herein, we profiled the serum glycomes of experimental mouse sepsis models to identify changes induced by 4 different clinical bacterial pathogens (Gram-positive: *Streptococcus pneumoniae, Staphylococcus aureus*, Gram-negative: *Escherichia coli* and *Salmonella* Typhimurium) using our lectin microarray technology. We observed global shifts in the blood sera glycome that were conserved across all four species, regardless of whether they were Gram positive or negative. Bisecting GlcNAc was decreased upon sepsis and a strong increase in core 1/3 *O*-glycans was observed. Lectin blot analysis revealed a high molecular weight protein induced in sepsis by all four bacteria as the major cause of the core 1/3 *O*-glycan shift. While the identity of this protein remains to be elucidated, its presence indicates a common feature of bacterial sepsis associated with this glycomic signature.

## Introduction

Sepsis is classified as an infection of the bloodstream that is associated with pathological inflammation and organ system dysfunction^1^. Currently, few diagnostic biomarkers and effective targeted therapies exist^1, 2^. A better understanding of the host-response and pathogenic mechanisms associated with disease onset and progression are required for developing more effective treatment strategies. Both Gram-positive and Gram-negative bacteria have been identified as common pathogens in septic patients. Incidence of sepsis driven by bacteria has increased in recent years, associated with the development of antibiotic resistance ^3, 4^. Bacteria initiate an innate immune response through host recognition of pathogen-associated molecular patterns (PAMPs). Examples of bacterial PAMPs include lipoteichoic acid (a component of the cell wall of gram-positive bacteria) and lipopolysaccharide (expressed by gram-negative bacteria) as well as constituents expressed by both gram-positive and gram-negative bacteria (peptidoglycan). These PAMPs initiate an inflammatory response through recognition by Toll-like receptors (TLRs). Uncontrolled stimulation of TLRs results in excessive inflammation associated with the septic phenotype ^5^.

Glycosylation is known to play a role in immunity and inflammation. Innate immune lectins can recognize glycan structures on the surface of bacteria and signal an immune response. Inflammatory cytokines can shift cell surface *N*-glycosylation of endothelial cells, contributing to inflammatory vascular diseases ^6, 7^. Recent work demonstrated that shifts in the glycan structure of a single glycoprotein in the context of discrete pathogens could help drive a septic state specific to Gram-positive bacteria in a TLR4-dependnent manner^8^. However, whether such glycomic shifts are observable at the global level is unknown.

Herein, we examined the global glycosylation profile of blood sera in mouse models of experimental sepsis. Previous studies have indicated that mouse models closely mimic human responses in inflammatory diseases^9^. We focused our work on sepsis induced by four different clinical isolates of bacterial pathogens: Gram-positive *Streptococcus pneumoniae* (SPN), methicillin-resistant *Staphylococcus aureus* (MRSA), Gram negative *Escherichia coli (*EC), and *Salmonella enterica* Typhimurium (ST). Sera from control as well as early and late sepsis time points were analyzed using our dual-color lectin microarray technology ^10^. This technology has been used to perform glycomics on a wide variety of samples including exosomes ^11, 12^, cervical lavage samples^13^ and human tumor tissues^14, 15^. Our glycomic analysis revealed two conserved changes occurring upon sepsis triggered by both Gram-positive and Gram-negative bacteria, a loss of bisecting GlcNAc and a dramatic increase in core 1/3 *O*-glycans. Lectin blot analysis confirmed our findings and indicated that a single protein may be responsible for the increased core 1/3 O-glycan signature. Overall, our work points towards a common mechanism for bacterially-induced sepsis marked by conserved changes in the glycome.

## Experimental Methods

### Laboratory Animals

Animal studies were performed by the Marth laboratory at the University of California Santa Barbara. All studies were done with the approval of the Institutional Animal Care and Use Committees of the University of California Santa Barbara and the Sanford-Burnham-Prebys Medical Discovery Institute. Animal experiments were carried out with adult 8–12-week-old mice with equal numbers of male and female mice. Mice were provided sterile pellet food and water *ad libitum*. Littermates of four or five animals per cage were housed in a pathogen-free barrier facility at the University of California Santa Barbara.

### Bacterial Strains and Culture Conditions

*Escherichia coli (EC)* strain ATCC 25922 (clinical isolate, FDA strain Seattle 1946), *Salmonella* enterica serovar Typhimurium (*ST*) reference strain ATCC 14028 (CDC 6516-60), *Streptococcus pneumoniae* (*SPN)* serotype 2 strain D39 and methicillin-resistant *Staphylococcus aureus* (*MRSA*) strain CA-MRSA USA300 were all used in these experiments^16–20^.

### Bacterial Infections

All mice were infected and monitored as previously described^8^ according to LD50 values that were pre-determined for each bacterial species by identifying the dose at which 50% of infected animals died post infection by the specified delivery routes. *EC* and *SPN* bacterial strains were administered as an intraperitoneal (i.p.) infection with an EC dose at 10X LD_50_ and *SPN* dose at 10X LD_50_. Gastric intubation was used for the administration of *ST* at a dose of 20X LD_50_, or i.p. at a dose 20X LD_50_. Intravenous infection was utilized for *MRSA* infection at a dose of 20X LD_50_. Blood was collected and the bacterial *cf.*u were measured at designated times post-infection. Mice that met minimal thresholds were utilized in this study and those outside of the target range were not analyzed further.

### Lectin Microarray

Samples were labeled with Alexa Fluor 555-NHS. Serum protein concentrations were determined using the DC assay. 50 μg of protein were labeled for each individual sample following the manufacturers protocol. Reference samples were created for each bacterial experiment and labeled with Alexa Fluor 647-NHS. For SPN, MRSA and ST, a bacteria-specific reference sample was prepared by mixing equal amounts of sera from all 48 animals used in each study. For EC, a master reference was created from the sera samples from the EC, ST and SPN experiments. Printing, hybridization and data analysis were performed as previously described ^10, 13, 21, 22^. The printlists for our lectin microarrays are shown in Supplemental Table 1. For each lectin microarray experiment, only lectins that were active on >30% of all arrays were considered in the analysis.

### Lectin blots

Sera samples were depleted of albumin prior to analysis by lectin blot using the CaptureSelect™ MultiSpecies Albumin Depletion Product (Thermo Fisher). Briefly, 100 μL of beads were used to deplete 5 μL of each serum sample. The concentration was then measured by DC assay and 20 μg of protein was resolved on an SDS page gel (gradient 4%-20%, BioRad) and transferred to a nitrocellulose membrane. Protein loading was visualized using Ponceau staining. Membranes were then blocked in 5% BSA in PBST (pH 7.4, 0.05% Tween-20) followed by washing with PBST (3x, 5 min).

Biotinylated MPL (5 μg/mL, Vector Laboratories) was then added to the membrane in blocking buffer (1 hr, r.t.) followed by washing with PBST (pH 7.4, 0.05% Tween-20, 3x, 5 min). Membranes were then incubated with Avidin-HRP (1:1000 in blocking buffer, ThermoFisher, 1 hr, r.t.) followed by three washes with PBST. Blots were developed with SuperSignal West Pico (ThermoScientific).

## Results and Discussion

### Description of experimental sepsis models

Although glycosylation plays important roles in infection and host-response, there is little known about overall glycomic changes in sera in response to bacterial sepsis. Recent studies on the glycosylation of specific glycoproteins in mouse models found associations between glycosylation and different mechanisms of sepsis caused by Gram-positive and Gram-negative bacteria ^8, 18, 23^. However, whether this translates into broader glycomic alterations in sera due to sepsis was not explored. To address this issue, we analyzed sera from a previously published experimental sepsis study ^8^. In this study, mice were infected with clinical isolates of bacterial strains that commonly induce sepsis: *Methicillin-resistant Staphylococcus aureus* (MRSA), *Streptococcus pneumoniae* (SPN), Escherichia *coli* (EC), and *Salmonella enterica* Typhimurium (ST). For each pathogen early and late post-infection timepoints (early and late sepsis), corresponding to colony forming units (cfu) thresholds, were analyzed (**Scheme 1**). A total of 48 mice were studied per bacterial species, with equal numbers of female and male mice (n=8 per sex per group: uninfected, pre-septic, septic, n=16 total for each condition).

### Lectin microarray analysis of glycomic response to bacterial sepsis from Gram-positive bacteria

The Gram-positive species *Streptococcus pneumoniae* (SPN) and Methicillin-resistant *Staphylococcus aureus* (MRSA) have both been named as priority pathogens by the World Health Organization because of their high burden of disease and antibiotic resistance ^24^. Both of these bacteria are important causes of human sepsis. *Streptococcus pneumoniae* (SPN) commonly colonizes mucosal surfaces of the human upper respiratory tract, and is a major cause of community-acquired pneumonia ^25^. Methicillin-resistant *Staphylococcus aureus* (MRSA) is an antibiotic resistant variant of a bacteria commonly found on the skin and in the upper respiratory tract. MRSA is a leading cause of bacterial infections in health-care and community settings. To look for glycomic signatures associated with bacterial sepsis caused by these organisms, we analyzed sera samples from our mouse models using our dual-color lectin microarray technology (**Scheme 2**) ^10, 21^.

Lectin microarrays display immobilized carbohydrate binding proteins with known glycan specificities to detect glycomic variations between samples ^10, 13, 21, 22^. In brief, sera samples (sample) and a bacteria-specific pooled reference (reference) were labeled with orthogonal fluorescent dyes. Equal amounts of protein (7 μg each) of sample and reference were analyzed on the lectin microarray (> 100 lectins, see **Table S1** for printlist). Only lectins passing our quality control are shown. Heatmaps displaying the normalized data for Gram-positive bacterial species (SPN and MRSA) are shown in **Figure 1** and **Supplementary Figures S1 & S2**.

**Figure 1:**
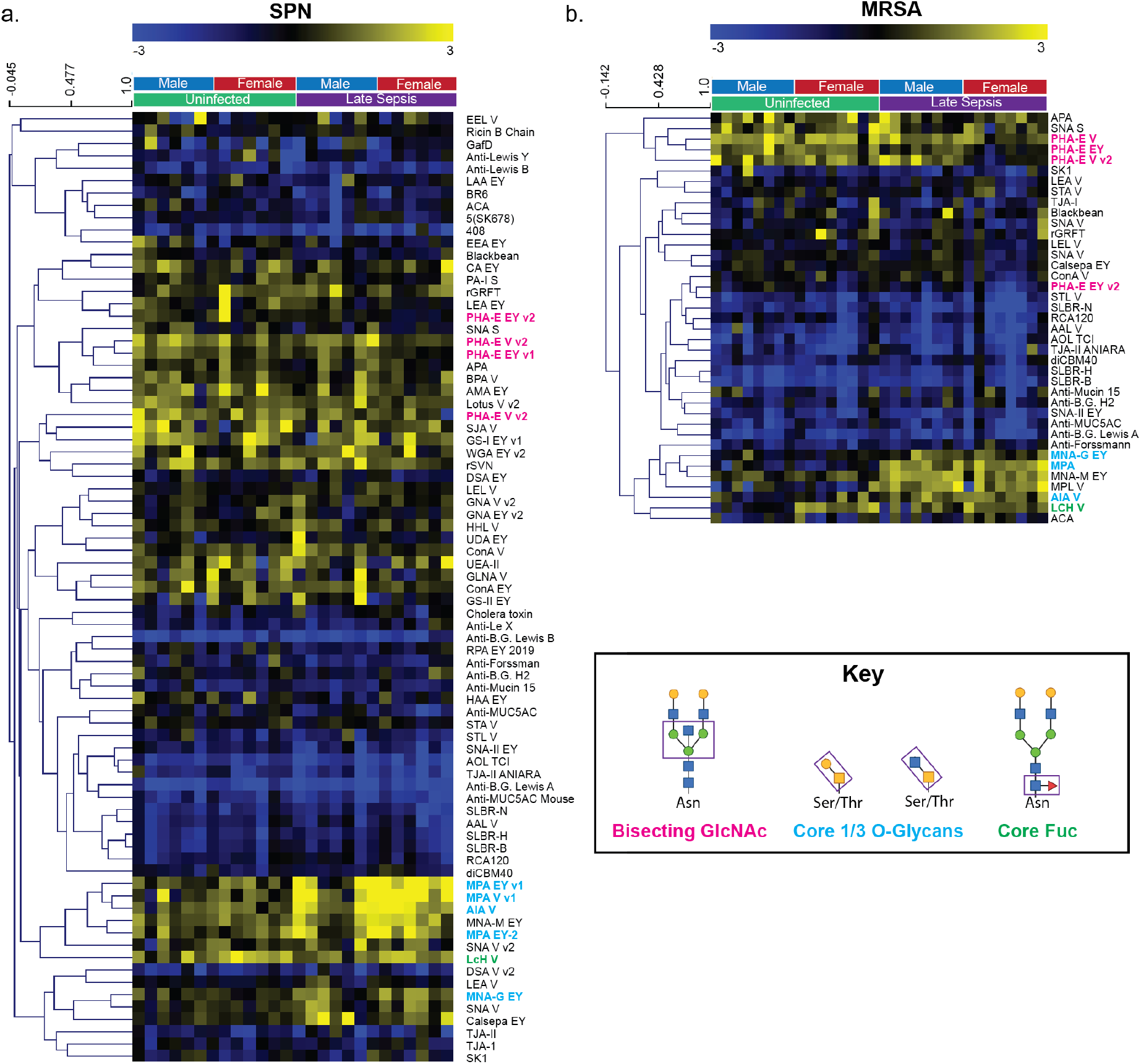
Heat map of lectin microarray data for Gram-positive bacteria. Median normalized log_2_ ratios (Sample (S)/Reference (R)) of mouse sera samples were ordered by uninfected or late sepsis for (a.) SPN and (b.) MRSA. Yellow, log_2_(S) > log_2_(R); blue log_2_(R) > log_2_(S). Lectins associated with bisecting GlcNAc (pink), Core 1/3 O-glycans (blue), and core fucose (green) are highlighted.

Although SPN and MRSA have different colonization patterns and tropism, we observed a remarkably consistent glycomic response by the host to sepsis induced by both pathogens (**Figure 1, Supplementary Figures S3 & S4**). The most striking change was a significant increase in core 1/3 *O*-Glycans (lectins: MPA, AIA, MNA-G, SPN: ~2-3-fold increase, *p* = 0.0007, MRSA: ~5-fold increase, *p =* 2 x 10^-8^, increase based on MPA). This observation is discussed in more detail below. We also observed a decrease in bisecting GlcNAc, which was more dramatic in the MRSA-infected animals (PHA-E, SPN:~ 1.5-fold decrease, *p* = 0.1, MRSA: ~1.5-fold decrease, *p =* 0.0001). The reduction of bisecting GlcNAc was detected at both early and late sepsis timepoints, indicating that this change occurs early in progression of sepsis (**Supplementary Figure S5**). Although the meaning of this change is unclear, it is of note that bisecting GlcNAc has a known role in IgG biology and is found on ~10% of all human IgG ^26, 27^. When on the Fc region of IgG, this glycan epitope increases affinity for the Fcγ3a receptor, leading to enhanced antibody dependent cellular cytotoxicity (ADCC). During sepsis, a loss of IgG correlates with an increase in mortality ^28^. Whether the reduction of this glycan structure is due to altered IgG levels or other glycoproteins remains to be examined.

### Lectin microarray analysis of glycomic response to bacterial sepsis from Gram-negative bacteria

The severity of inflammation in sepsis induced by Gram-negative bacteria has been shown to be higher than that of Gram-positive bacteria 29. The gut pathogens *Salmonella enterica* Typhimurium (ST) and *Escherichia coli* (EC) are common causes of sepsis in clinic 30, 31. To explore the glycomic response to Gram-negative bacterial sepsis, we performed lectin microarray analysis on samples from ST and EC infected mice as previously described. Heatmaps are shown in Figures 2 & 3 and **Supplementary Figures S6 & S7**.

**Figure 2:**
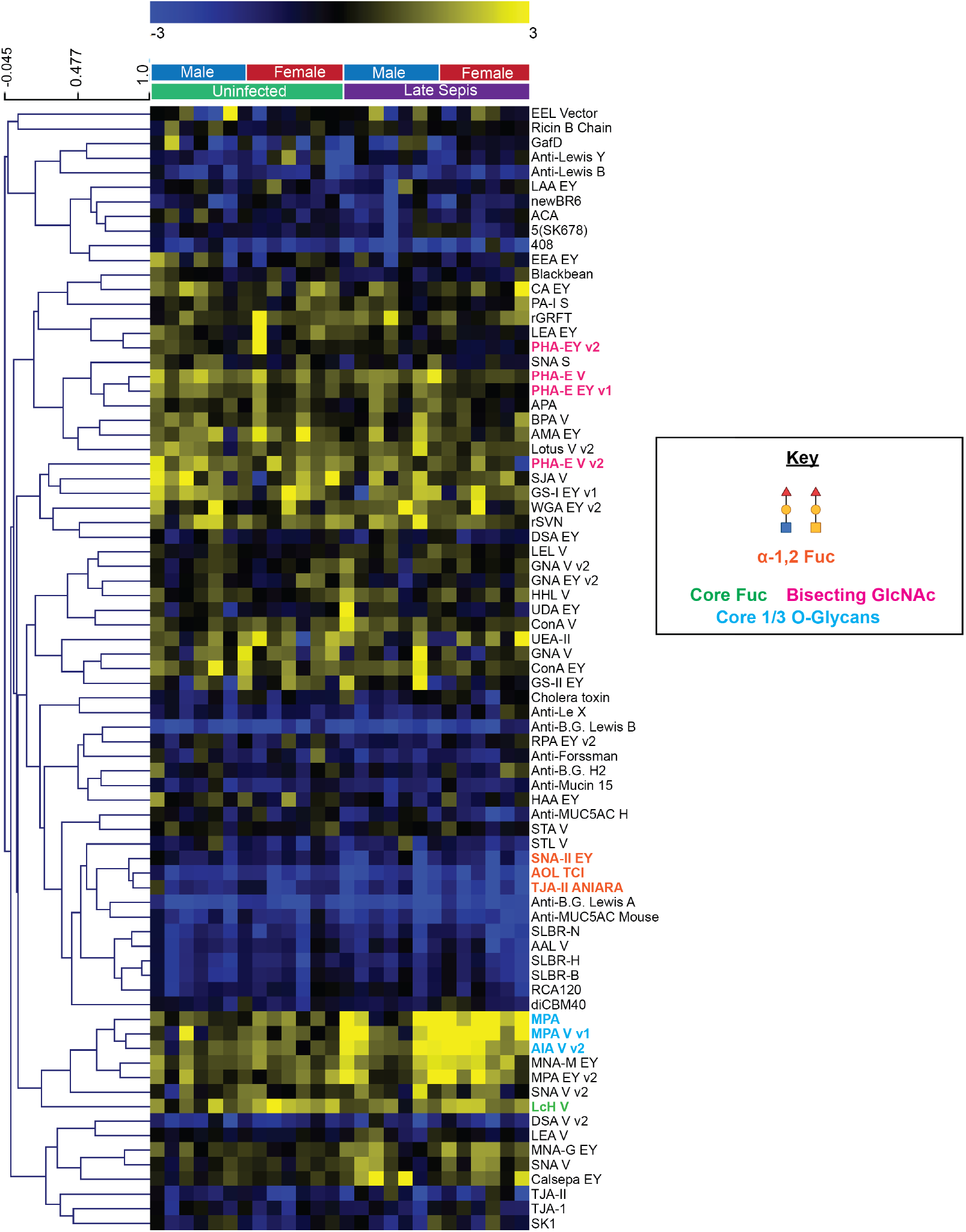
Heat map of lectin microarray data for sera from ST infected mice. Median normalized log_2_ ratios (Sample (S)/Reference (R)) of mouse sera samples were ordered by uninfected or late sepsis for (a.) SPN and (b.) MRSA. Yellow, log_2_(S) > log_2_(R); blue log_2_(R) > log_2_(S). Lectins associated with bisecting GlcNAc (pink), Core 1/3 O-glycans (blue), core fucose (green) and α-1,2 fucose (orange) are highlighted.

**Figure 3:**
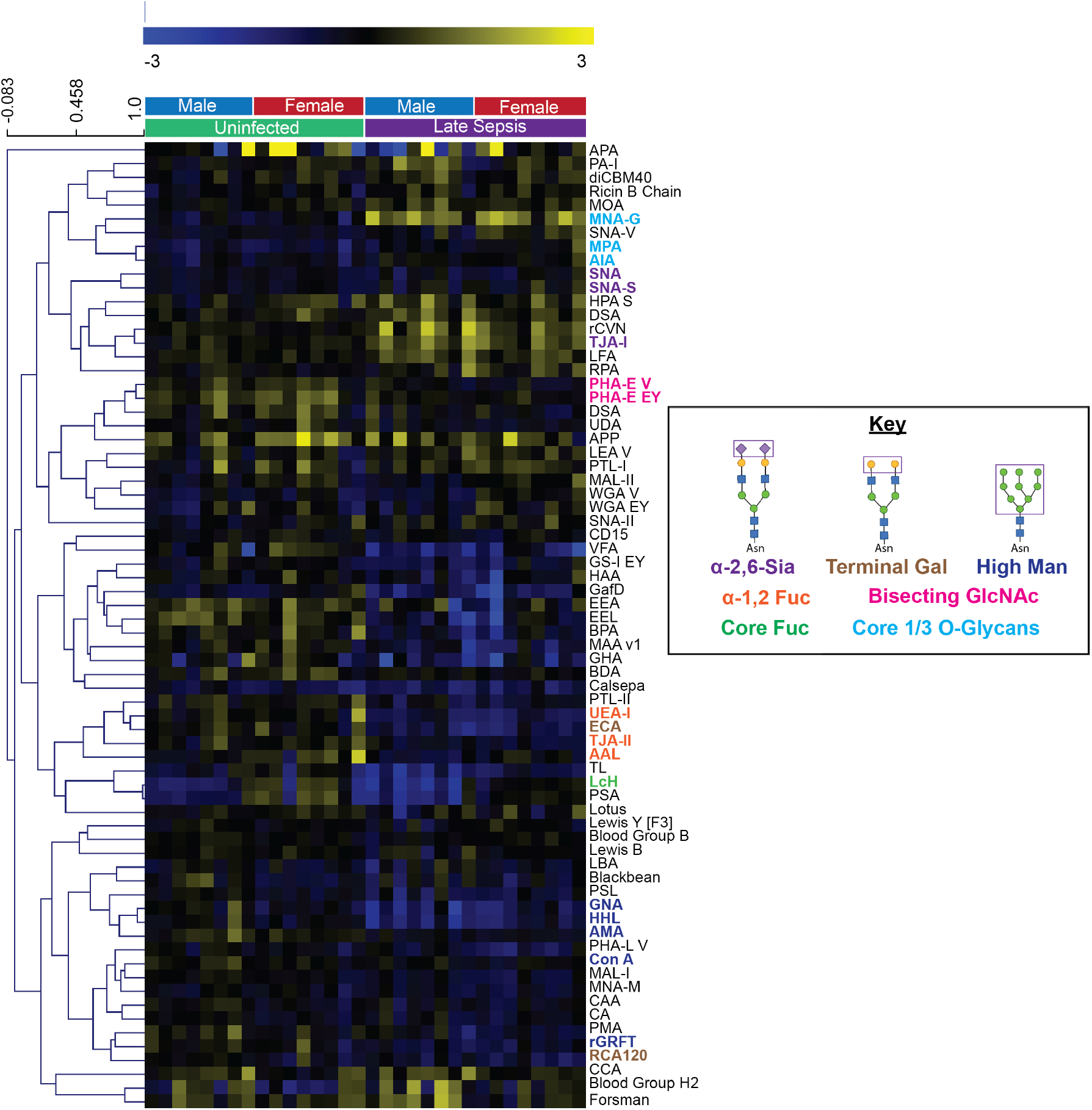
Heat map of lectin microarray data for sera from EC infected mice. Median normalized log_2_ ratios (Sample (S)/Reference (R)) of mouse sera samples were ordered by uninfected or late sepsis for (a.) SPN and (b.) MRSA. Yellow, log_2_(S) > log_2_(R); blue log_2_(R) > log_2_(S). Lectins associated with bisecting GlcNAc (pink), Core 1/3 O-glycans (blue), core fucose (green), α-1,2 fucose (orange) α-2,6 sialic acid (purple), terminal galactose (brown), and high mannose (blue) are highlighted.

We observed conserved responses between Gram-negative and Gram-positive bacteria (**Figures 2 & 3, Supplementary Figures S9 & S10**). Again, one of the strongest changes observed in sepsis for both Gram-negative bacterial species was the increase in core 1/3 *O*-Glycans (MPA, AIA, MNA-G, ST: ~4-fold increase, *p =* 0.01, EC: ~1.5-fold increase, *p =* 0.0001). A decrease in bisecting GlcNAc epitopes was also observed (PHA-E, ST: ~1.5-fold decrease, *p =* 0.03, EC: ~1.5-fold decrease, *p =* 0.0001). In EC, but not ST, the response of core 1/3 *O*-Glycans and bisecting GlcNAc to infection could be clearly seen in early sepsis (**Supplementary Figure S8**). Of the four bacteria studied, ST has the longest incubation period, due to its oral route of infection. Unlike the other bacteria where the early and late timepoints are 24 h apart, the two timepoints in ST are collected 3 days apart (**Scheme 1**). This may explain the delayed response observed.

Bacterial sepsis from ST and EC showed both overlapping and unique glycan signatures. Both Gram-negative bacteria induced a loss of α1,2-fucosylation (ST: ~ 2-fold decrease, lectins: TJA-II, SNA-II, AOL, EC: ~1.5-fold decrease, lectins: UEA-I, AAL, PTL-II). In the sera, α1,2-fucosylation is controlled by FUT2. This enzyme has a powerful role in the establishment and maintenance of the gut microbiota, which these two pathogens may affect. In recent work, α1,2-fucosylation has been found to increase colonization of *Salmonella* Typhimurium in a mouse model ^32^. Conversely, the lack of FUT2, and thus sera α1,2-fucosylated glycans, is associated with an increase in severity for enteric EC infections ^33^. The response to sepsis caused by EC was distinguishable from ST by additional changes in the serum glycome. EC-induced sepsis correlated with a loss of high- and oligomannose structures (GRFT, HHL, ConA, AMA, GNA), an increase in α2,6-sialic acids (SNA. TJA-I), a concomitant decrease in terminal β-galactose (RCA, ECA) and an increase in sulfated or α2,3-sialylated glycans (MAA, MAL-I). These differences were unique to sepsis caused by this bacterial pathogen.

### Increase in core 1/3 *O*-glycans is conserved in sepsis across bacterial species and may indicate a common host-response

All four bacterial species studied cause a striking increase in core 1/3 *O*-glycan levels at late stage sepsis, when signs of sepsis are visually overt (**Figures 1–3 and 4a**. This effect on core 1/3 *O*-glycans is seen in both Gram-positive and Gram-negative infections. The increase of core 1/3 *O*-glycans can be observed in early sepsis in three out of the four organisms studied. The one exception is *Salmonella* Typhimurium, which may again be due to the longer incubation time for sepsis in this oral infection model.

**Figure 4:**
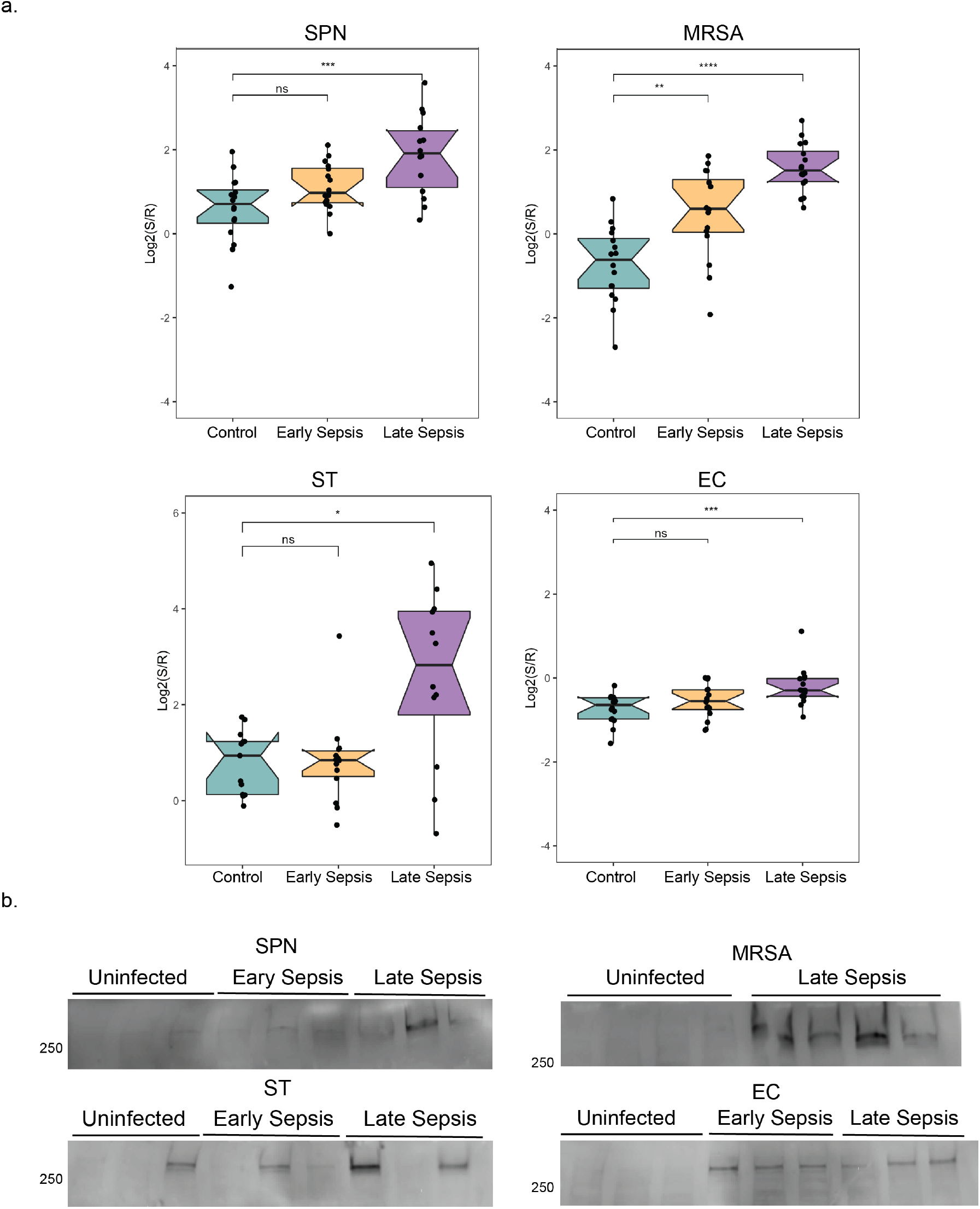
Core 1/3 O-glycan levels increase during sepsis. a. Box plot analysis of lectin binding by MPA (core 1/3 O-glycans) are depicted. *P*-values derive from Student’s t-test (* *p* < 0.05, ** < 0.01, *** < 0.001, **** < 0.0001). b. Lectin blots probed with biotinylated MPA and Streptavidin HRP show an increase in a single high molecular weight band in both early and late sepsis. Whole blots and corresponding Ponceau stained membranes are shown in **Figure S11**.

Core 1 *O*-glycans have been observed on several sera-derived glycoproteins. These include IgA and IgD immunoglobulins and the proteoglycan, lubricin ^34, 35^. To corroborate our lectin microarray analysis, we performed lectin blot analysis of mouse sera samples from our experimental models using the core 1/3 *O*-glycan binding lectin MPA. Gels were run using standard reducing conditions, prior to transfer. Interestingly, we do not observe a general increase in core 1/3 *O*-glycans across multiple glycoproteins in sepsis. Instead in all four experimental models of bacterial sepsis, we observe what appears to be a single, high molecular weight band (>250 kD, **Figure 4b**, **Supplementary Figure S11**). The staining of this band correlated with the observed MPA signal in our lectin microarrays. For example, in both our lectin blot and lectin microarray data, we see a variable response in ST-induced sepsis. In cases where the lectin microarray shows high levels of MPA, we observe high levels of the high MW glycoprotein in the MPA lectin blot and vice versa. This indicates that the dramatic change in sera core 1/3 *O*-glycosylation levels observed may be due to a single protein, the identity of which remains to be elucidated.

## Conclusions

Bacterial sepsis is one of the top 10 causes of human death and disability, and its frequency is increasing with antibiotic resistance ^3, 4^. The Gram-positive (MRSA and SPN) and gram-negative (EC and ST) bacterial pathogens used in our study are clinical isolates and common causes of sepsis. At the molecular level, sepsis is incompletely defined, with a descriptive rather than molecular diagnosis. Currently, there are few diagnostic biomarkers and with the exception of early application of antibiotics, targeted therapy has been ineffective ^1, 2^. There is a need to study further the molecular underpinnings of this syndrome. Glycosylation is known to play roles in immunity and inflammation, but the roles of glycans in sepsis are not well defined. In this current study, we compared the sera glycomes of uninfected and septic mice across multiple bacterial strains (MRSA, SPN, EC and ST) at two post-infection timepoints, tethered to increasing blood cfu levels ^8^.

We observed major changes in the sera glycome that were conserved across all four experimental sepsis models, regardless of whether bacterial pathogens were Gram positive or negative. We observed a common decrease in bisecting GlcNac and an increase in Core 1/3 *O-*glycans associated with bacterial sepsis caused by multiple pathogens. The identification of proteins bearing these glycan changes may ultimately identify a common pathogenic pathway, similar to that achieved in detecting the altered glycans of AP isoenzymes in the pathogenesis of Gram-negative sepsis^8^. These changes were visible at the earliest post-infection times studied. Lectin blot analysis identified what appears to be a conserved high molecular weight protein with elevated levels of core 1/3 *O*-glycans in sepsis caused by all bacteria studied. While the identity of this protein remains to be elucidated, its presence indicates a common mechanism in bacterial sepsis associated with this glycomic signature. Whether this protein is a novel glycoprotein secreted in response to sepsis, or a new glycoform of a circulating glycoprotein that has been remodeled in response to sepsis ^36^ remains to be established. Future studies will be required to identify this mechanism and see whether it is applicable to blood samples from human sepsis patients.

## Acknowledgements

Research undertaken herein by J.D.M. and P.V.Z. was supported by NIH grants HL131474 and DK048247. This research was also supported by funding from the Canada Excellence Research Chairs Program (L. Mahal).

---

**Scheme 1:**
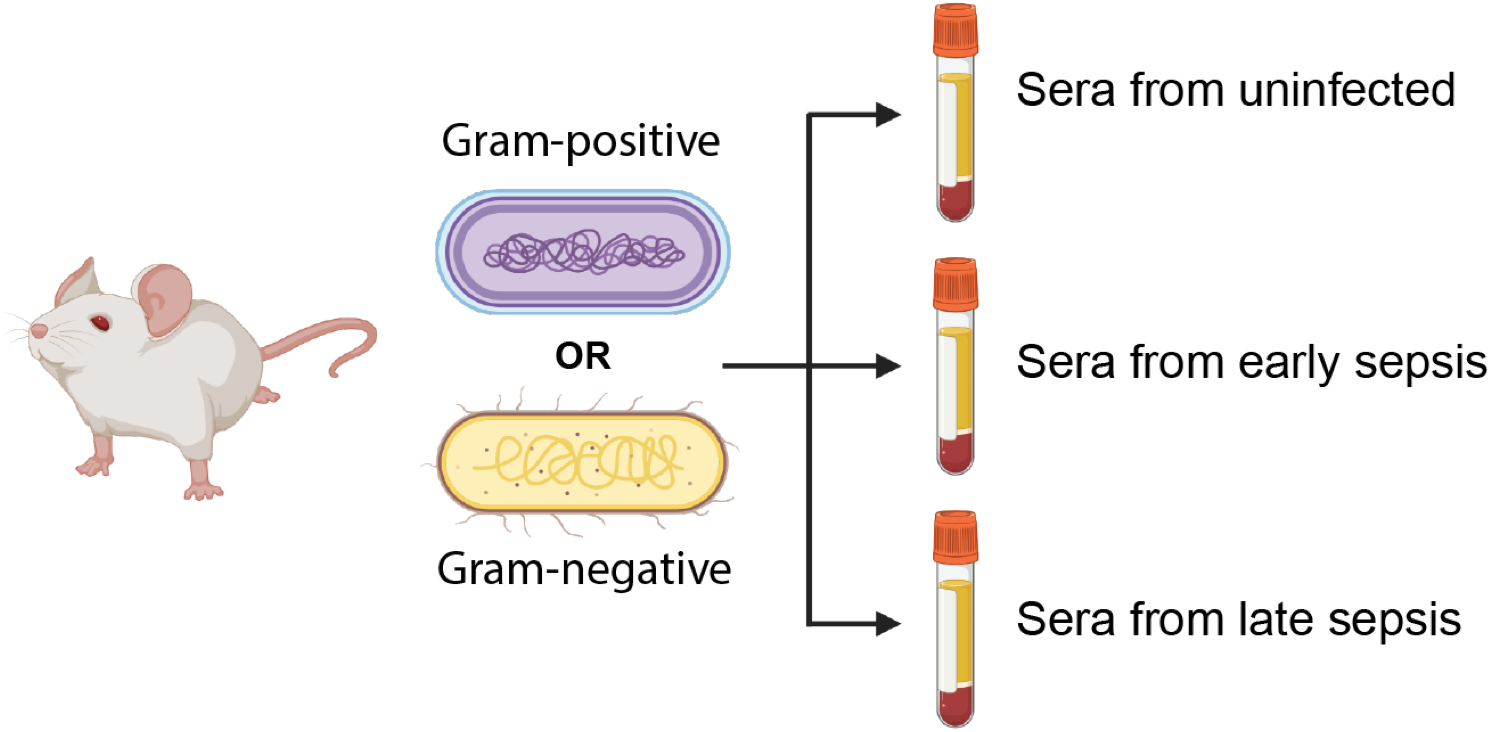
Murine model of sepsis. Sera was collected from uninfected mice and all comparisons were made to this group for each type of bacteria. Blood was collected at specified times post infection to determine bacterial cf.u for both early and late stages of sepsis as described ^8^.

**Scheme 2:**
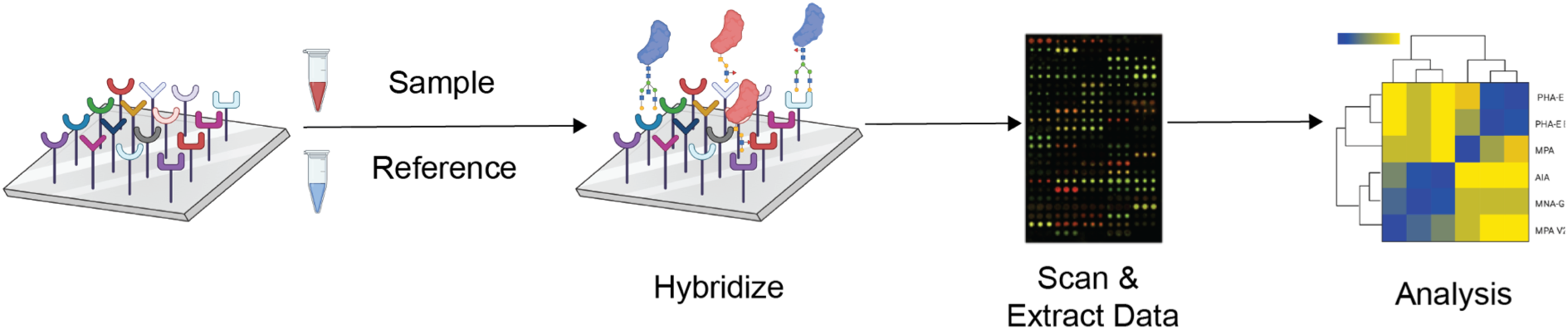
Lectin microarray workflow. Equal protein amounts (7 μg) for each sample and an orthogonally labeled mixed reference were combined and analyzed on the lectin microarray.

