## Supplemental Figure for "Glycomic analysis reveals a conserved response to bacterial sepsis induced by different bacterial pathogens"

Table of Contents

**Supplementary Figure S1**. Heat map of lectin microarray data for sera from SPN infected

**Supplementary Figure S2.** Heat map of lectin microarray data for sera from MRSA infected mice

**Supplementary Figure S3.** Volcano plot depicting changes in glycan abundances between septic and control animals infected with SPN

**Supplementary Figure S4.** Volcano plot depicting changes in glycan abundances between septic and control animals infected with MRSA.

**Supplementary Figure S5.** Bisecting GlcNAc levels decrease during sepsis (SPN and MRSA)

**Supplementary Figure S6.** Heat map of lectin microarray data for sera from ST infected mice

**Supplementary Figure S7.** Heat map of lectin microarray data for sera from EC infected mice

**Supplementary Figure S8.** Bisecting GlcNAc levels decrease during upon sepsis induced by EC and SPN

**Supplementary Figure S9.** Volcano plot depicting changes in glycan abundances between septic and control animals infected with ST

**Supplementary Figure S10.** Volcano plot depicting changes in glycan abundances between septic and control animals infected with EC

**Supplementary Figure S11.** MPA Lectin Blots and Corresponding Ponceaus

**Supplemental Table 1.** Lectins used in microarrays

**Supplemental Table 2:** Lectin Microarray Information

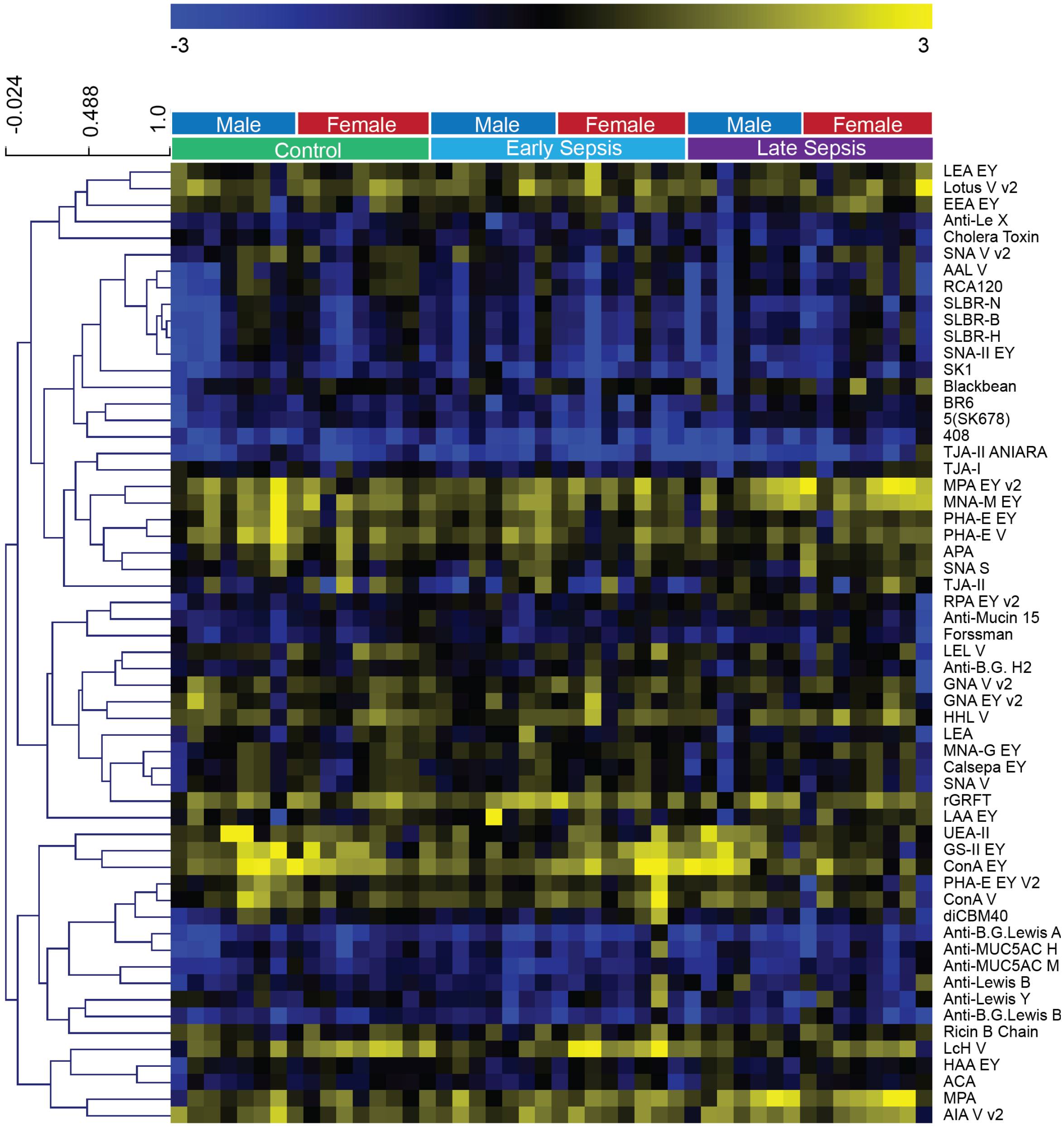


**Supplementary Figure S1: Heat map of lectin microarray data for sera from SPN infected mice** Median normalized $\log_2$ ratios (Sample (S)/Reference (R)) of mouse sera samples were ordered by uninfected, early , and late sepsis.

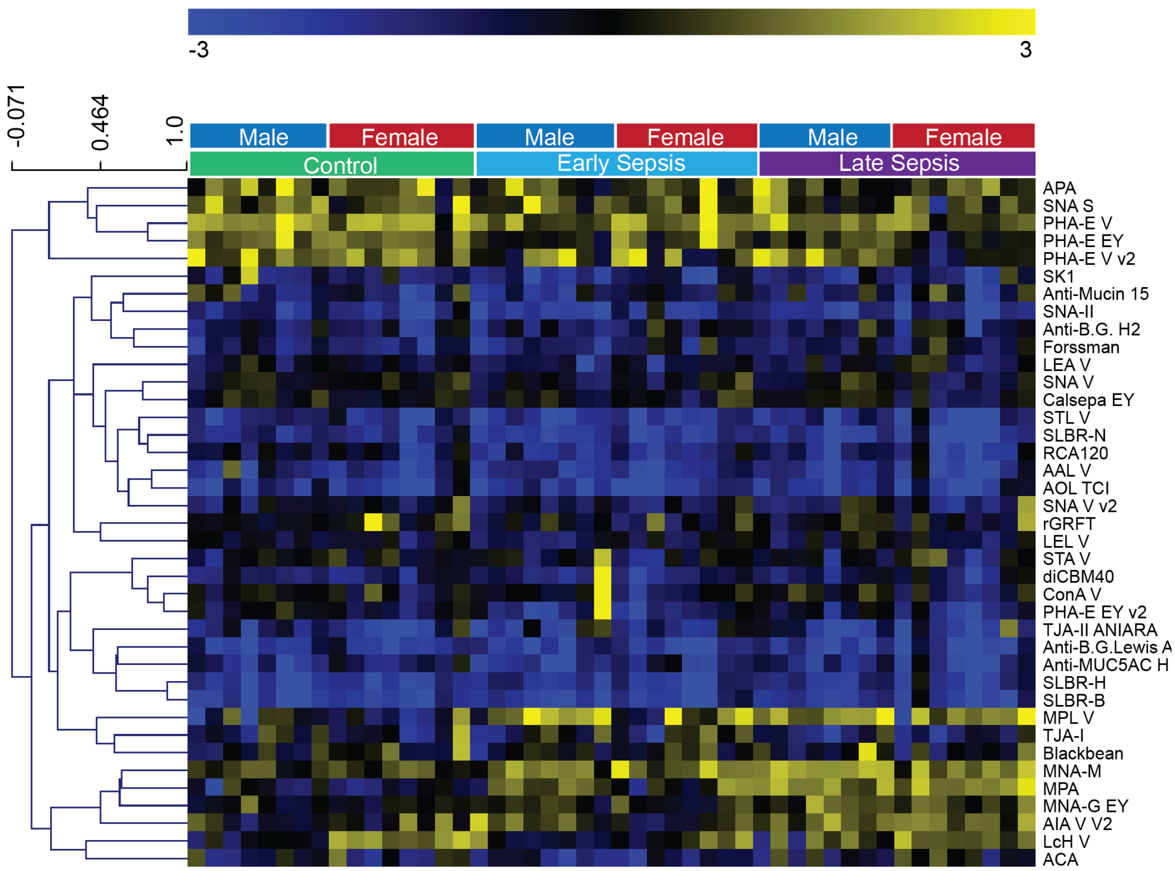


**Supplementary Figure S2: Heat map of lectin microarray data for sera from MRSA infected mice** Median normalized $\log_2$ ratios (Sample (S)/Reference (R)) of mouse sera samples were ordered by uninfected, early, and late sepsis.

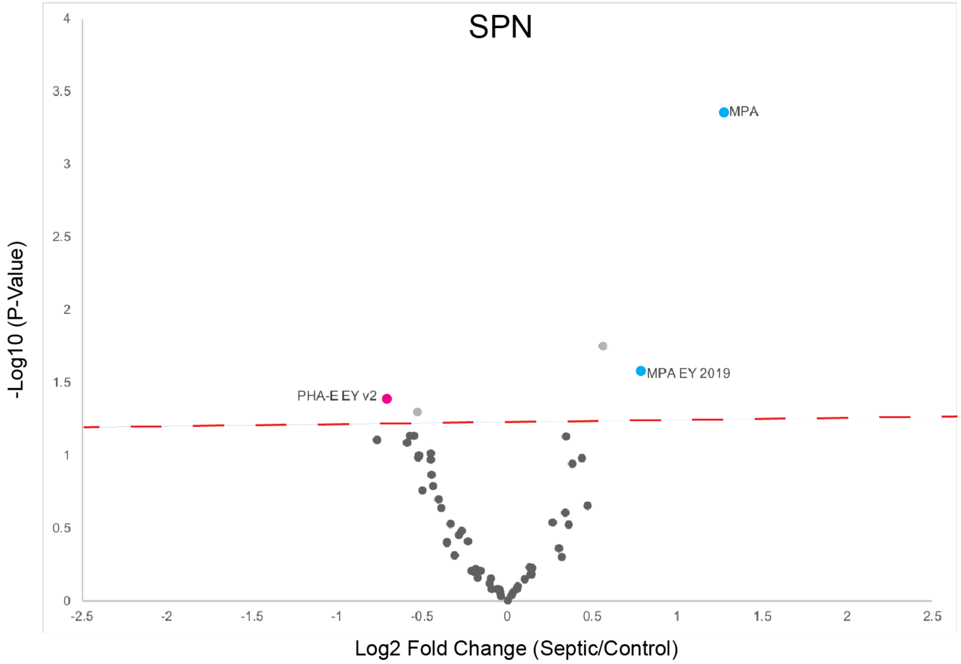


**Supplementary Figure S3: Volcano plot depicting changes in glycan abundances between septic and control animals infected with SPN** Statistically significant lectins specific for core 1/3 *O*-glycans ( blue) and bisecting GlcNAc (pink) are labeled. The dashed red line indicates $p < 0.05$ by the Students t-test.

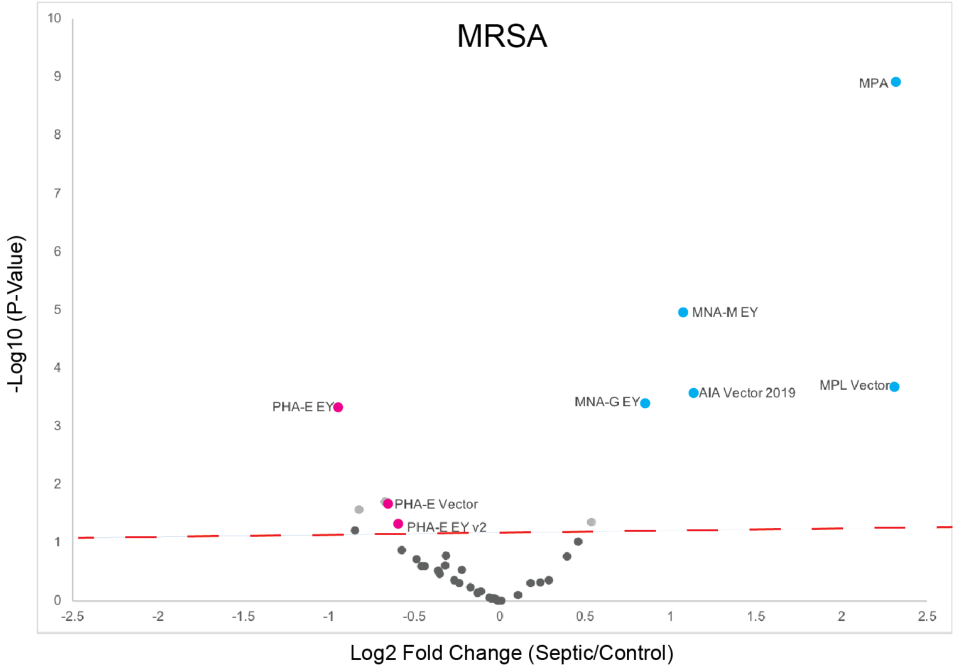


**Supplementary Figure S4: Volcano plot depicting changes in glycan abundances between septic and control animals infected with MRSA.** Statistically significant lectins specific for core 1/3 *O*-glycans ( blue) and bisecting GlcNAc (pink) are labeled. The dashed red line indicates $p < 0.05$ by the Students t-test.

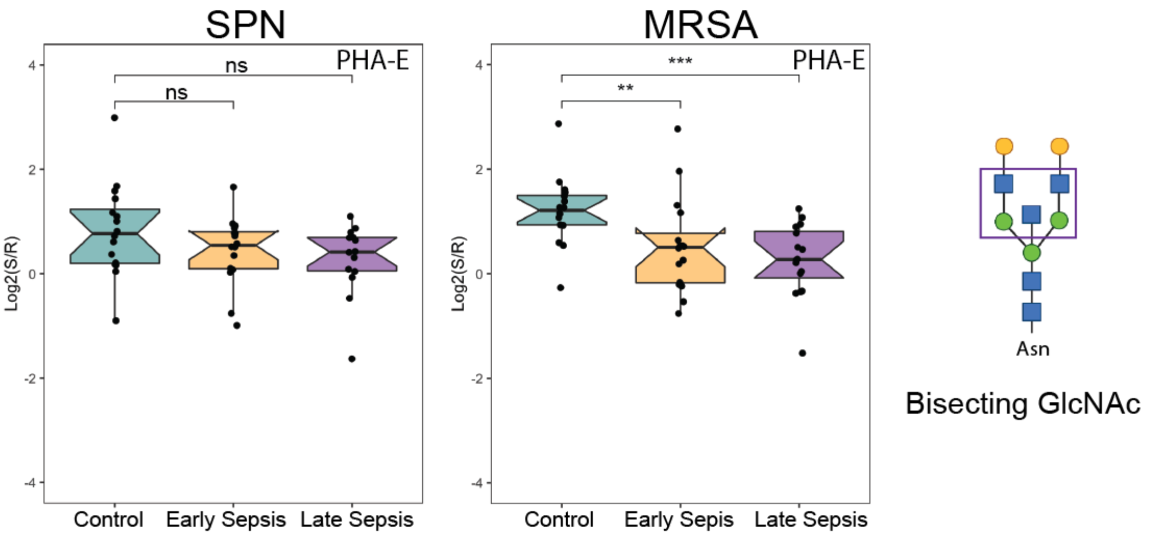


**Supplementary Figure S5: Bisecting GlcNAc levels decrease during sepsis (SPN and MRSA)** Box plot analysis of bisecting GlcNAc lectin binding by PHA-E. *P*-values derive from Student's t-test (* *p* ≤ *0.05,* ** ≤ *0.01*, *** ≤ 0.001).

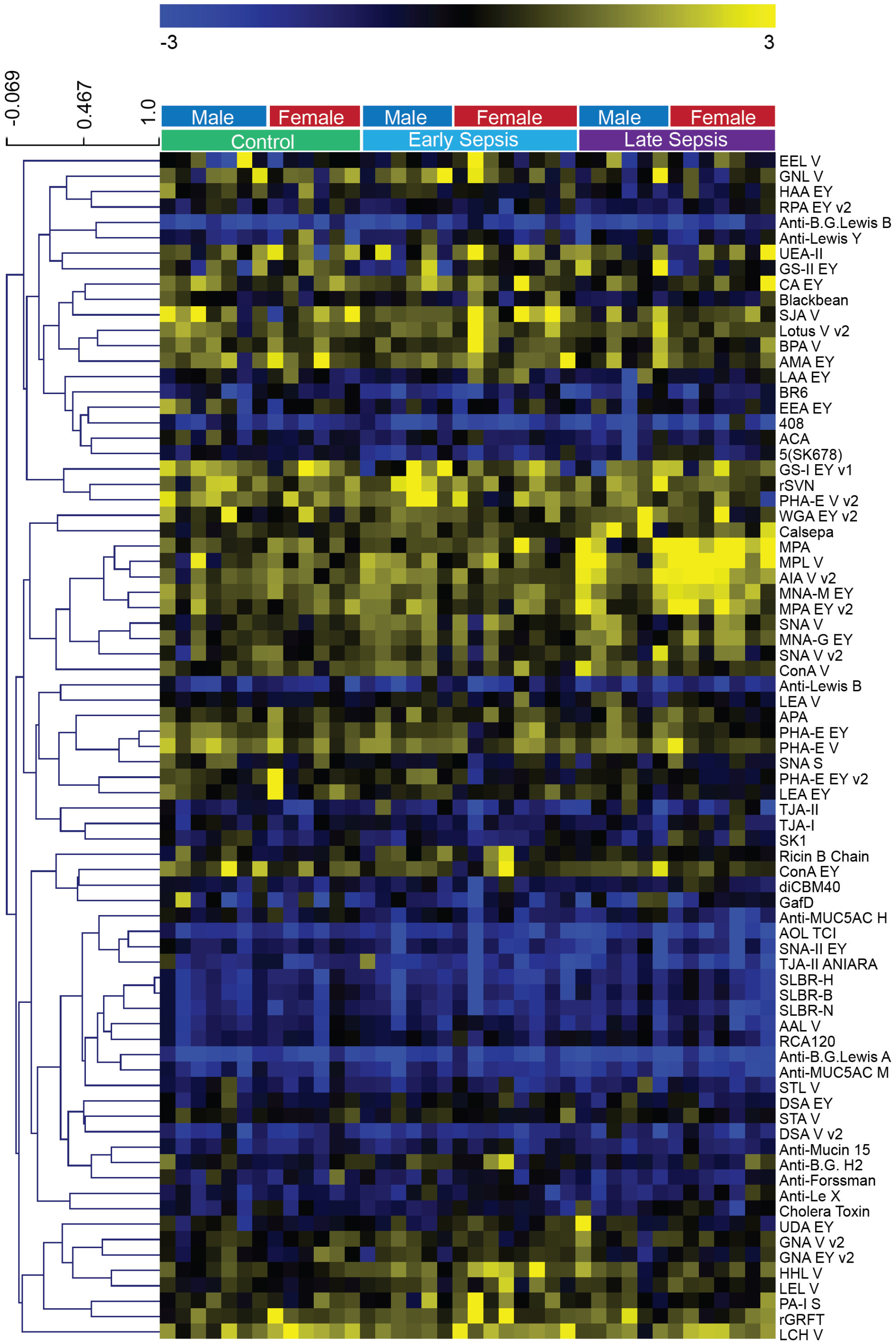


**Supplementary Figure S6: Heat map of lectin microarray data for sera from ST infected mice** Median normalized $\log_2$ ratios (Sample (S)/Reference (R)) of mouse sera samples were ordered by uninfected, early, and late sepsis.

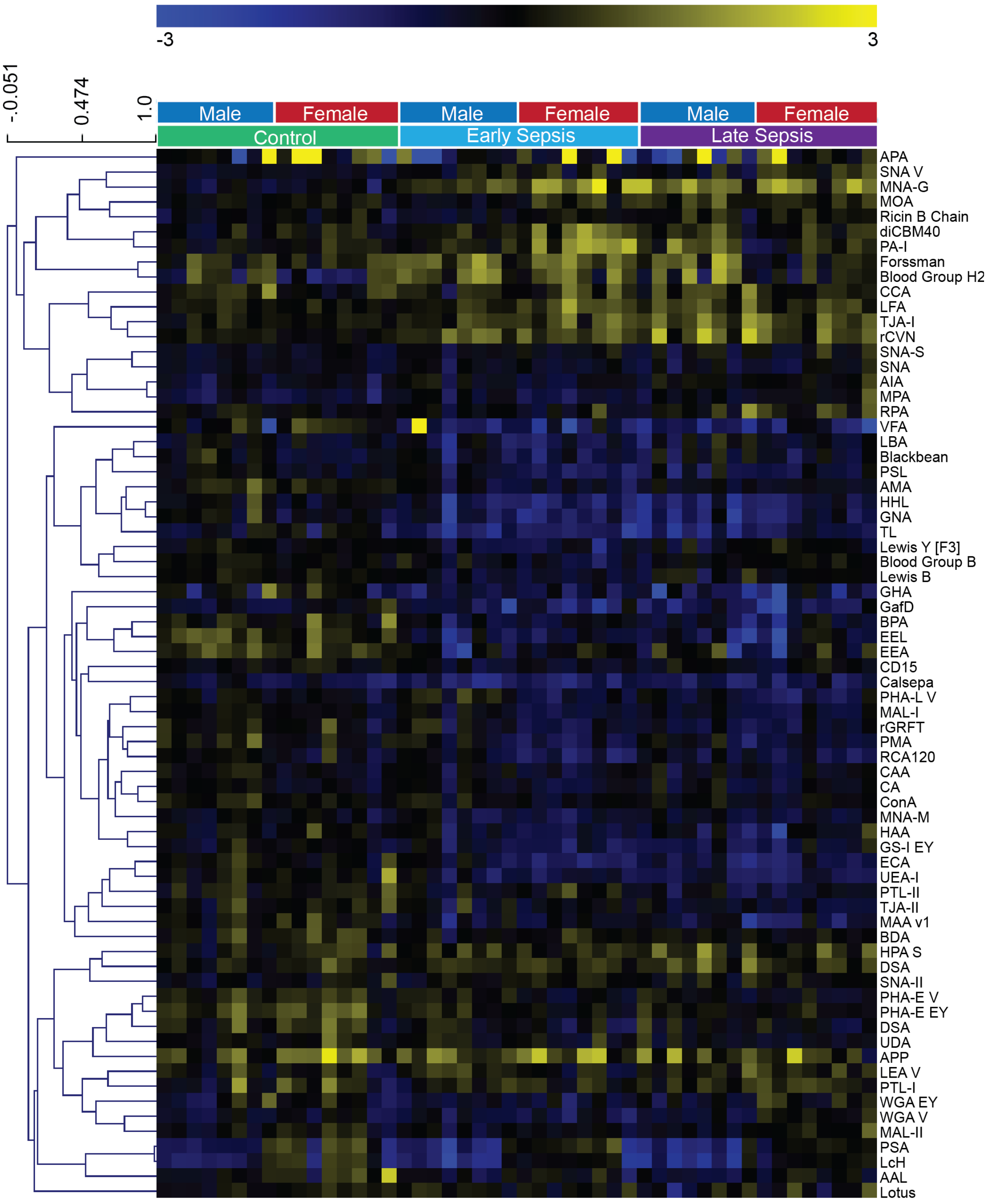


**Supplementary Figure S7: Heat map of lectin microarray data for sera from EC infected mice** Median normalized $log_2$ ratios (Sample (S)/Reference (R)) of mouse sera samples were ordered by uninfected, early, and late sepsis.

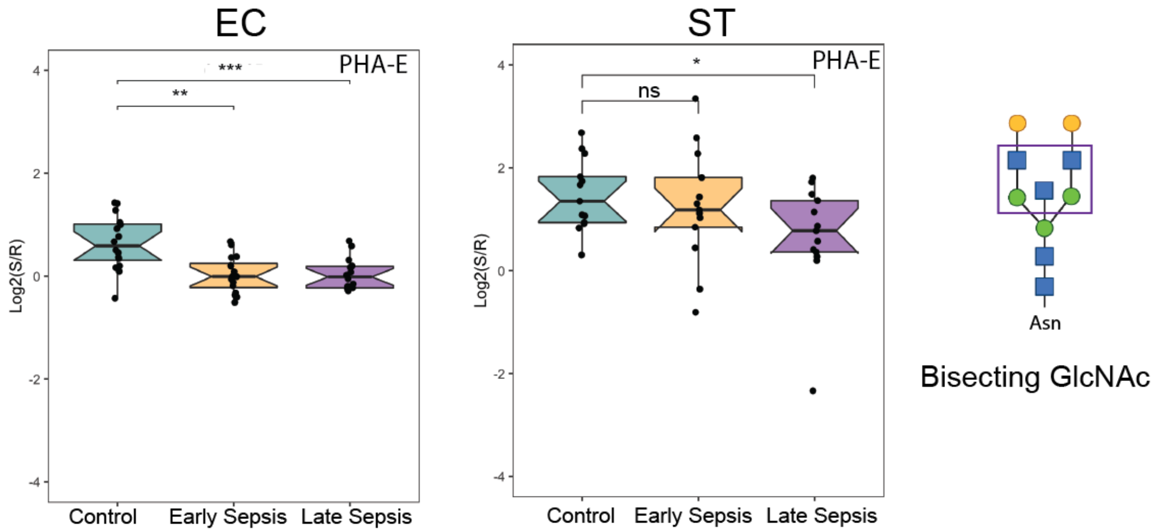


**Supplementary Figure S8: Bisecting GlcNAc levels decrease during upon sepsis induced by EC and SPN** Box plot analysis of bisecting GlcNAc lectin binding by PHA-E. *P*-values derive from Student's t-test (* $p \leq 0.05$, ** $\leq 0.01$, *** $\leq$ 0.001).

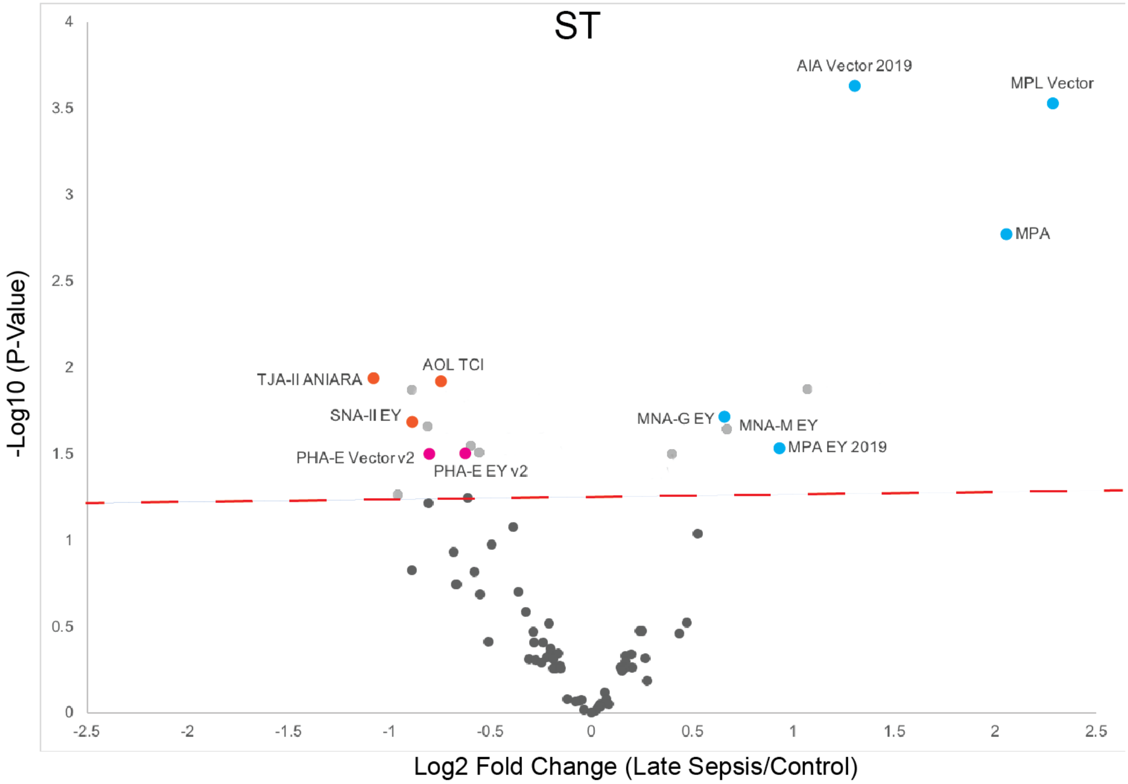


**Supplementary Figure S9: Volcano plot depicting changes in glycan abundances between septic and control animals infected with ST** Statistically significant lectins specific for core 1/3 *O*-glycans ( blue), bisecting GlcNAc (pink) and α-1,2 fucose (orange) are labeled. The dashed red line indicates $p < 0.05$ by the Student's t-test.

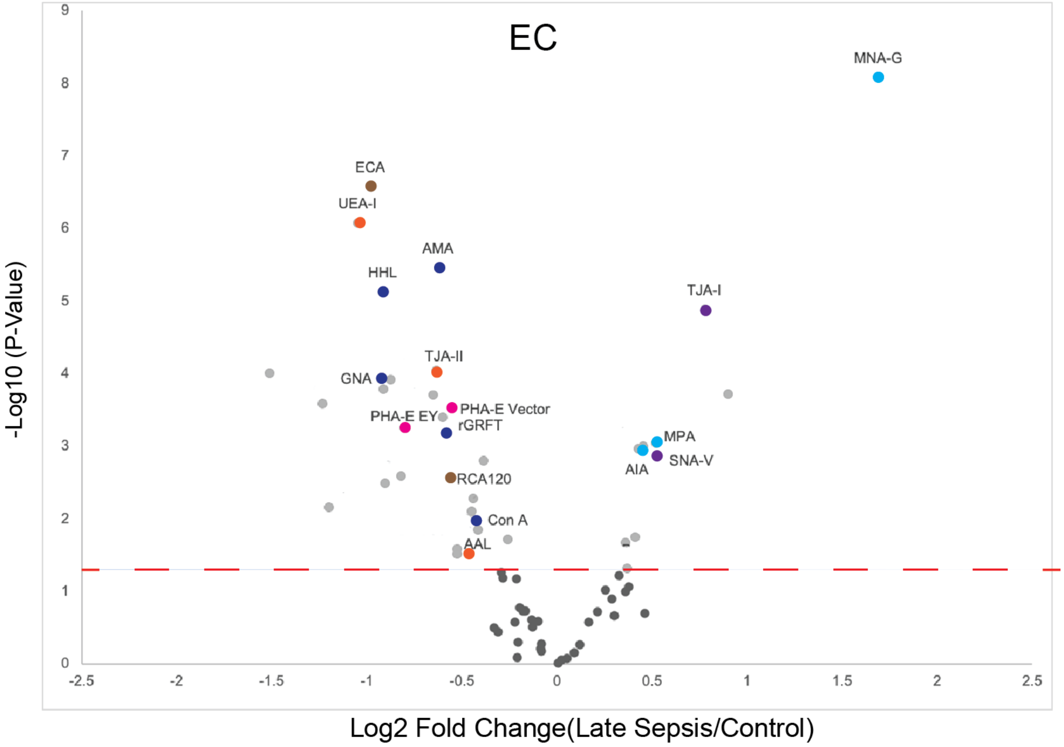


**Supplementary Figure S10: Volcano plot depicting changes in glycan abundances between septic and control animals infected with EC.** Statistically significant lectins specific for core 1/3 *O*-glycans ( blue), bisecting GlcNAc (pink), α-1,2 fucose (orange), terminal galctose (brown), high mannose (dark blue), and α-2,6 sialic acid (purple) are labeled. The dashed red line indicates $p < 0.05$ by the Student's t-test.

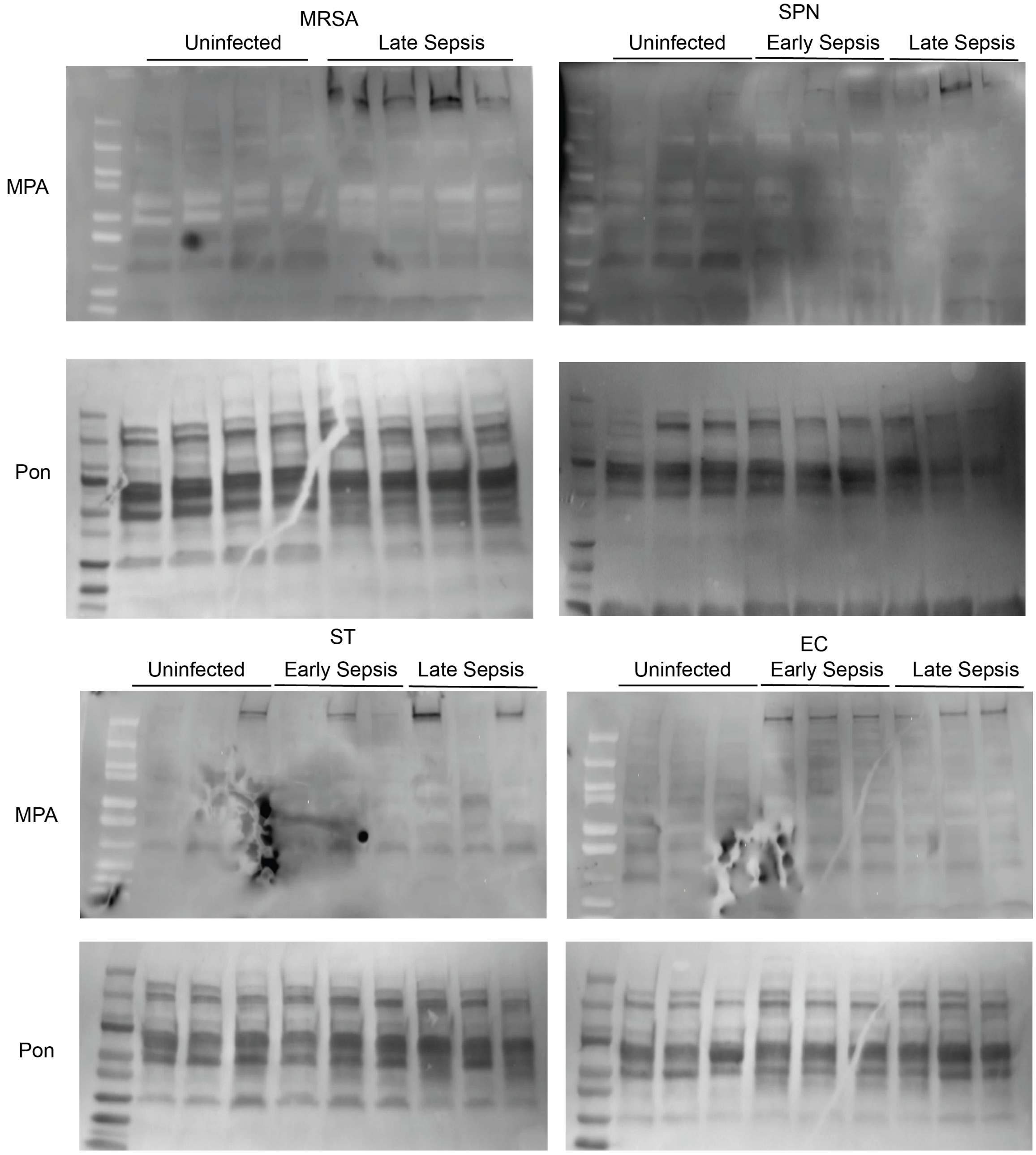


**Supplementary Figure S11: MPA Lectin Blots and Corresponding Ponceaus.** The entire lectin blots for data shown in Figure 5 are shown along with Ponceau analysis that confirms even loading.

### Supplemental Table 1: Lectins used in microarrays

| Lectin | Species/Origin | Print Conc. (µg/mL) | Rough Specificity /Inhibitory monosaccharide | Vendor/Source |
|---|---|---|---|---|
| AAL [1] | *Aleuria aurantia* | 1000 | Fucose | Vector |
| ACA [1] | *Amaranthus Caudatus* | 1000 | Gal-β1,3-GalNAc | Vector |
| AIA [1] | *Artocarpus integrifolia* | 500 | β1,3-GalNAc | Vector/EY |
| AMA [1] | *Allium moly* | 500 | Oligo mannose | EY |
| Anti-B.G.H2 [1] | MAb mouse IgM [A46-B/B10] | undiluted | Blood group H2 antigen | Santa Cruz Biotechnology |
| Anti-CD15 [1] | MAb mouse IgM [MY-1] | undiluted | Lewis X | Abcam |
| Anti-Forssman [1] | MAb Rat IgM [117C9] | undiluted | Forssman Antigen | Abcam |
| Anti-Lewis B | IgM [T218] | undiluted | Lewis B | Sigma |
| Anti-Lewis X [1] | MAb mouse IgM [P12] | undiluted | Lewis X | Abcam |
| Anti-Lewis Y [1] | MAb mouse IgM [F3] | undiluted | Lewis Y | Abcam |
| Anti-MUC5AC human [1] | Mab mouse IgG1 [CLH2] | undiluted | human MUC5AC | Sigma |
| Anti-MUC5AC mouse | Goat polyclonal to mouse MUC5AC | undiluted | mouse MUC5AC | LSBio |
| Anti-Mucin 15 [1] | Mab mouse IgG1 [H-5] | undiluted | Mucin 15 | Santa Cruz Biotechnology |
| Anti-Sialyl Lewis A [1] | Mab mouse IgG1 | undiluted | Sialyl Lewis A | Abcam |
| Anti-Sialyl Lewis X [1] | Mab mouse IgM | undiluted | Sialyl Lewis X | Abcam |
| AOL [1] | *Aspergillus oryzae* | 1000 | Fucose | TCI America |
| APA [1] | *Abrus precatorius* | 500 | Gal-β1,3-GalNAc / Lac | EY |
| ASA [1] | *Allium sativum* | 1000 | Mannose | EY |
| Blackbean [1] | *Blackbean crude* | 1000 | GalNAc | EY |
| BPA [1] | *Bauhinia purpurea* | 500 | β-Gal / β-GalNAc | Vector |
| BR6 | unknown (from unpublished work) | 480 | under investigation | Gift from Dr. Barbara Bensing |
| CA | *Colchicum autumnale* | 1200 | Bi-antennary N-linked glycans | EY |
| CAA [1] | *Caragana arborescens* | 1000 | Bi-antennary N-linked glycans | EY |
| Calsepa [1] | *Calystegia sepium* | 1000 | Bisecting N-linked glycans | EY |
| CCA [1] | *Cancer antennarius* | 1000 | 9-O-Acetly sialylation / 4-O-Acetyl sialylation | EY |
| Cholera Toxin [1] | *Vibrio cholerae* | 1000 | GM1 ganglioside | Sigma |

| | | | | |
|---|---|---|---|---|
| Con A | *Canavalia ensiformis* | 1000 | Tri-mannose core | EY/Vector |
| CSA [1] | *Cystisus scoparius* | 1000 | Terminal GalNAc | EY |
| DBA [1] | *Dolichos Biflorus* | 1000 | GalNAc | Vector |
| diCBM40 [1] | engineered NanI from *Clostridium perfringens* | 1000 | α Sialylation | Generated in house |
| DSA [1] | *Datura stramonium* | 500 | LacNAc | EY/Vector |
| ECA [1] | *Erythrina cristagalli* | 1000 | LacNAc | Vector |
| EEL/EEA [1] | *Eunonymus europaeus* | 1000 | Blood Group B | Vector/EY |
| GafD [1] | recombinant GafD from *Escherichia coli* | 1000 | GlcNAc | Generated in house |
| GHA [1] | *Glechoma hederacea* | 500 | GalNac | EY |
| GNA/GNL [1] | *Galanthus nivalis* | 1500 | Oligo mannose | Vector/EY |
| GS-I [1] | *Griffonia simplicifoia-I* | 1000 | α-Gal / Lac | Vector/EY |
| GS-II [1] | *Griffonia simplicifoia-II* | 1000 | GlcNAc | Vector |
| GS-IB4 [1] | *Griffonia simplicifoia-I, isolectin B4* | 2000 | Gal | Vector |
| H84T [1] | *Banana lectin* | 1000 | High mannose | Gift from Dr. David Markovitz |
| HAA [1] | *Homarus americanus* | 1000 | Terminal GalNAc | EY |
| HHL [1] | *Hippeastrum Hybrid* | 1500 | Oligo/High mannose | Vector |
| HPA [1] | *Helix pomatia* | 1000 | Blood Group A | Sigma/EY |
| IRA [1] | *Iris Hybrid* | 1000 | GalNAc / Lac | EY |
| LAA | *Laburnum alpinum* | 900 | GlcNAc | EY |
| LBA [1] | *Phaseolus lunatus* | 1000 | Blood Group A | EY |
| LcH [1] | *Lens Culinaris* | 1000 | Core Fucose | Vector |
| LEA/LEL [1] | *Lycopersicon esculentum* | 1000 | GlcNAc | Vector/EY |
| LFA [1] | *Limax flavus* | 500 | α Sialylation | EY |
| Lotus [1] | *Lotus tetragonolobus* | 1000 | Fucose | Vector |
| MAA [1] | *Maackia amurensis* | 500 | Sialylation/Sulfatio n | EY |
| MAL-I [1] | *Maackia amurensis-I* | 2000 | Sialylation/Sulfatio n | Vector |
| MAL-II [1] | *Maackia amurensis-II* | 2000 | Sialylation/Sulfatio n | Vector |

| | | | | |
|---|---|---|---|---|
| MNA-G [1] | *Morus nigra Morniga G* | 1000 | GalNAc | EY |
| MNA-M [1] | *Morus nigra Morniga M* | 1000 | Oligo mannose / Gal | EY |
| MPA/MPL [1] | *Maclura pomifera* | 1000 | β1,3-GalNAc | Vector |
| NPA [1] | *Narcissus pseudonarcissus* | 1000 | Oligo mannose | Vector |
| PA-I | *Pseudomonas aeruginosa* | 1000 | Gal | Sigma |
| PA-IL [1] | *bacteria* | 1000 | GalNAc | Generated in house |
| PHA-E [1] | *Phaseolus vulgaris Erythroagglutinin* | 1000 | Bisecting GlcNAc | Vector/EY/Sigma |
| PHA-L [1] | *Phaseolus vulgaris Leukoagglutinin* | 1000 | β1,6 Branching N-Link glycans | Vector/EY/Roche |
| PMA [1] | *Polygonatum multiflorum* | 500 | Oligo mannose | EY |
| PNA [1] | *Arachis hyogaea* | 1000 | Gal-β1,3-GalNAc | Vector/EY |
| PSA [1] | *Pisum sativum* | 1000 | Core Fucose | Vector |
| PSL [1] | *Polyporus squamosus* | 1000 | α2,6 sialylation | EY |
| PTA [1] | *Psophocarpus tetragonolobus* | 500 | Blood Groups | EY |
| PTL-I [1] | *Psophocarpus tetragonolobus-I* | 1500 | Blood Group A | Vector |
| PTL-II [1] | *Psophocarpus tetragonolobus-II* | 1000 | α2 Fucose | Vector |
| RCA120 [1] | *Ricinus Communis Agglutinin I* | 1000 | Gal / Lac | Vector |
| rCVN [1] | *recombinant Cyanovirin* | 1000 | High mannose | Gift from Dr. Barry O'Keefe |
| rGRFT [1] | *recombinant Griffithsin* | 1000 | High mannose | Gift from Dr. Barry O'Keefe |
| Ricin B Chain [1] | *Ricinus communis* | 1000 | Gal | Vector |
| RPA [1] | *Robinia pseudoacacia* | 500 | Complex N-link glycans | EY |
| rSVN [1] | *recombinant Scytovirin* | 1000 | High mannose | Gift from Dr. Barry O'Keefe |
| SBA [1] | *Glycine max* | 1000 | LacdiNAc | Vector |
| SJA [1] | *Sophora japonica* | 1000 | LacdiNAc | Vector |
| SK1 | *Streptococcus sanguinis SK1* | 1800 | α2,3 sialylation | Gift from Dr. Barbara Bensing |
| SK678 | *Streptococcus sanguinis SK678* | 450 | α2,3 sialylation | Gift from Dr. Barbara Bensing |
| SLBR-B | *Streptococcus gordonii M99* | 1000 | α2,3 sialylation | Gift from Dr. Barbara Bensing |
| SLBR-H | *Streptococcus gordonii DL1* | 2000 | α2,3 sialylation | Gift from Dr. Barbara Bensing |
| SLBR-N | *Streptococcus gordonii UB10712* | 1000 | α2,3 sialylation | Gift from Dr. Barbara Bensing |
| SNA [1] | *Sambucus nigra* | 500/1000 | α2,6 sialylation | Vector/Sigma |
| SNA-II [1] | *Sambucus nigra-II* | 1000 | α2 Fucose /oligo mannose | EY |

| | | | | |
|---|---|---|---|---|
| STA/STL [1] | *Solanus tuberosum* | 500 | GlcNAc | Vector |
| TJA-I [1] | *Trichosanthes japonica-I* | 1000 | α2,6 sialylation | TCI |
| TJA-II [1] | *Trichosanthes japonica-II* | 1000 | α2 Fucose | NorthStar Bioproducts/Aniara Diagnostica |
| TL [1] | *Tulipa sp.* | 700 | GlcNAc | EY |
| UDA [1] | *Urtica dioica* | 1000 | GlcNAc / Oligo mannose | EY |
| UEA-I [1] | *Ulex europaaeus-I* | 1000 | α2 Fucose | Vector |
| UEA-II [1] | *Ulex europaaeus-II* | 2000 | GlcNAc | Vector |
| VFA [1] | *Vicia faba* | 1000 | GlcNAc | EY |
| VVA [1] | *Vicia villosa* | 1000 | Terminal GalNAc | Vector/EY |
| VVA(man) [1] | *Vicia villosa* | 500 | Mannose | Vector/EY |
| X408 | unknown (from unpublished work) | 1000 | under investigation | Gift from Dr. Barbara Bensing |
| WFA [1] | *Wisteria floribunda* | 1000 | GalNAc-β1,4 | Vector |
| WGA [1] | *Triticum vulgare* | 1000 | GlcNAc | Vector/EY |

[1]: lectins printed in the first set of lectin microarrays
[2]: lectins printed in the second set of lectin microarrays

### Supplemental Table 2: Lectin Microarray Information

| | **Description*** |
|---|---|
| **1. Sample: Glycan-containing sample (e.g. glycan, glycoprotein, cell lysate etc.)** | |
| Description of Sample | Sera was collected from uninfected mice as well as mice infected with different types of bacteria (EC, ST, MRSA, SPN). Sera was collected at two different time points for the infected mice that correspond to early and late sepsis that are defined by c.fu in the blood. All samples were collected in the Marth Laboratory at UCSD. |
| Sample preparation protocol | Sera samples were collected in the Marth Laboratory at UCSD and shipped to NYU and supplemented with protease inhibitor cocktails. |
| Labelling protocol for sample detection | Samples were labelled with Alexa Fluor 555-NHS (Thermo Fisher). Serum protein concentrations were determined using |

| | |
|---|---|
| | the DC assay. 50 $\mu$g of protein were labelled for each individual sample following the manufacturers protocol. |
| Two-color reference (if used) | Reference samples were created for each bacterial experiment and labeled with Alexa Fluor 647-NHS. For SPN, MRSA and ST, a bacteria-specific reference sample was prepared by mixing equal amounts of sera from all 48 animals used in each study. For EC, a master reference was created from the sera samples from the EC, ST and SPN experiments. |
| Assay protocol | Lectin microarrays are blocked with blocking buffer for one hour at room temperature. Slides are rinsed twice with PBST (0.005%) and once with PBS, then dry the slide using a slide spinner. Each slide was mounted on a 24-well format hybridization cassette (Arrayit), in which each well contains a subarray. To each well, add equal amounts of samples and universal reference, and dilute with PBS and PBST (0.2%) to reach the final volume (150uL). Incubate the slides on an orbital shaker for two hours at room temperature in the dark. After hybridization, wash the arrays with PBST (0.005%) twice for ten minutes, and twice for five minutes. Once finished, remove the slides from the cassette, and immerse the slides in ultrapure water, and dry the slides using a slide spinner. |
| **2. Lectin Library** | |
| General description of the lectin library used in the array | Lectin microarrays are generated in house. |
| List of lectins and glycan binding proteins, source, concentration and buffer | Please see **Supplemental Table 1.** |
| Modification of lectins (e.g. biotin) if any. | N/A |
| **3. Immobilization Surface; e.g., Microarray Slide** | |
| Immobilization surface | Nexterion Slide H Barcoded 3D Hydrogel Coated |
| Manufacturer | Shott North America |

| | |
|---|---|
| Custom preparation of surface | N/A |
| **4. Array Production** | |
| Description of Arrayer | Nano-Plotter 2.1 piezoelectric printer (GeSim, Germany) with cooled microwell plate holder and cooled printing deck |
| Lectin deposition | Three replicates of each lectin are printed onto each subarray. |
| Printing conditions | Dilute lectins to the pre-determined concentrations in the print buffer (final concentration of print buffer: 0.01% Tween-20, 1mM monosaccharide in PBS; Please see **Supplemental Table 1** for the concentrations of lectins). Load the mixed solution to the microplate. Before printing, check the humidity of the print chamber. The humidity should be kept around 50% during the entire printing. Ensure both microwell plate holder and printing deck are cooled. Adjust the cooling temperature based on ambient temperature and the temperature of the cooled slide deck surface, preventing moisture building up inside the print chamber. Once printing is complete, allow the slides to dry for at least one hour. |
| Array layout | For each microarray, it contains 24 subarrays (3 columns and 8 rows). In each subarray, triplicates of a lectin are printed, and for a row with five lectins, the spot layout should be 15 columns. The row number depends on how many lectin probes are printed on the arrays (i.e., 110 lectins require 22 rows). |
| Quality control | Well-characterized glycoproteins including fetuin, asialofetuin and RNase B are used for quality assurances of the printed microarrays. |
| **5. Detector and Data Processing** | |
| Instrument (scanner, flow cytometer) | Fluorescent Slide Scanner Genepix 4300A (Molecular Devices) |
| Instrument settings | Preview the slide to adjust photomultiplier gain (PMT) for each channel (Alexa Fluor-555: 532nm, Alexa Fluor-647: 635nm) so that the signals are not saturated and within the linear detection range. |
| Image analysis software | GenePix Pro 7 (Molecular Devices) |

| | |
|---|---|
| Data processing and statistical analysis | Extracted data is processed for quality checks using Grubbs outlier test with = 0.05. $Log_2$ values of the average signals are median-normalized over the individual subarray in each channel. |
| **6. Lectin Microarray Data Presentation** | |
| Data presentation and interpretation | Hierarchical clustering of the processed data is performed using Pearson Correlation coefficient, and visualized with Multi-experiment Viewer (MeV, v4.8, TM4 Microarray Software Suite). If a lectin's SNR (signal-to-noise ratio) < 3 for more than one third of the total samples, then this lectin is considered as inactive and excluded from the list. *P*-values are calculated using nonparametric statistical tests, which are generated by R (v3.6.1). |
| **7. Data Location** | |
| Data Location | |
